# Dissociating the Hallucinogenic and Neuroplastic Effects of Psilocybin

**DOI:** 10.64898/2026.04.06.716778

**Authors:** Jacob J. Baker, Emily Kogan, Shaorong Ma, Ju Lu, Yi Zuo

**Affiliations:** Department of Molecular, Cell and Developmental Biology, University of California Santa Cruz, Santa Cruz, CA 95064, USA; Department of Biological Sciences, Lehigh University, Bethlehem, PA 18015, USA

## Abstract

It is unclear how serotonin 2A receptors (5-HT_2A_Rs) in cortical layer 5 pyramidal neurons (L5 PyrNs) differentially contribute to psilocybin-induced hallucinations versus neuroplasticity. Here we show that psilocybin promotes synapse formation and maturation while accelerating the elimination of pre-existing synapses. Cell type-specific manipulation further demonstrated that 5-HT_2A_R signaling in L5 PyrNs is necessary and sufficient for psilocybin-induced synaptic remodeling but dispensable for the head-twitch response, a rodent proxy of hallucination.

## Main text

Psychedelics are psychoactive substances best known for their ability to induce hallucinations in humans^1^. In rodents, they acutely elicit the head-twitch response (HTR), a rapid, paroxysmal headshake well-established as a proxy for a serotonergic psychedelic’s hallucinogenic potential in humans^2^. Besides such psychotropic effects, psychedelics are potent promoters of neuroplasticity, elevating dendritogenesis and synapse formation while altering neurotransmission both *in vitro* and *in vivo*^3–5^. Psilocybin, a psychedelic derived from *Psilocybe* “magic” mushrooms, receives much research interest as a potential treatment for various neuropsychiatric disorders^6^; its neuroplastic effects are postulated to be pivotal to its sustained therapeutic value^5,7–9^. Like other serotonergic psychedelics, psilocybin is an agonist of the serotonin 2A receptor (5-HT_2A_R), which is abundantly expressed in the brain, particularly in neocortical layer 5 pyramidal neurons (L5 PyrNs)^10^. Pharmacological studies have implicated 5-HT_2A_R signaling in the hallucinogenic potential of psychedelics^2,11^, but its role in the neuroplastic effects remains controversial^12–17^. Recently, researchers have synthesized several plasticity-promoting, non-hallucinogenic psychedelic analogs^18–23^, but the role of L5 PyrNs in psilocybin’s neuroplastic versus hallucinogenic effects remains incompletely understood^17,24^.

To evaluate the hallucinogenic and neuroplastic effects of psilocybin in wild type (WT) mice, we administered psilocybin at doses of 0.3, 1, or 3 mg/kg bodyweight to ∼3 months old C57BL/6J mice. All doses induced robust HTRs; the 1 and 3 mg/kg doses elicited HTRs comparably, but significantly more than the 0.3 mg/kg dose (Fig. 1a). There is a modest sex difference in the 1.0 mg/kg psilocybin group, with females exhibiting slightly more HTRs than males (Extended Data Fig. 1a). HTR counts gradually increased during the first 5 min after injection, peaked between 5-10 min, and declined afterwards (Extended Data Fig. 1b).

**Figure 1.**
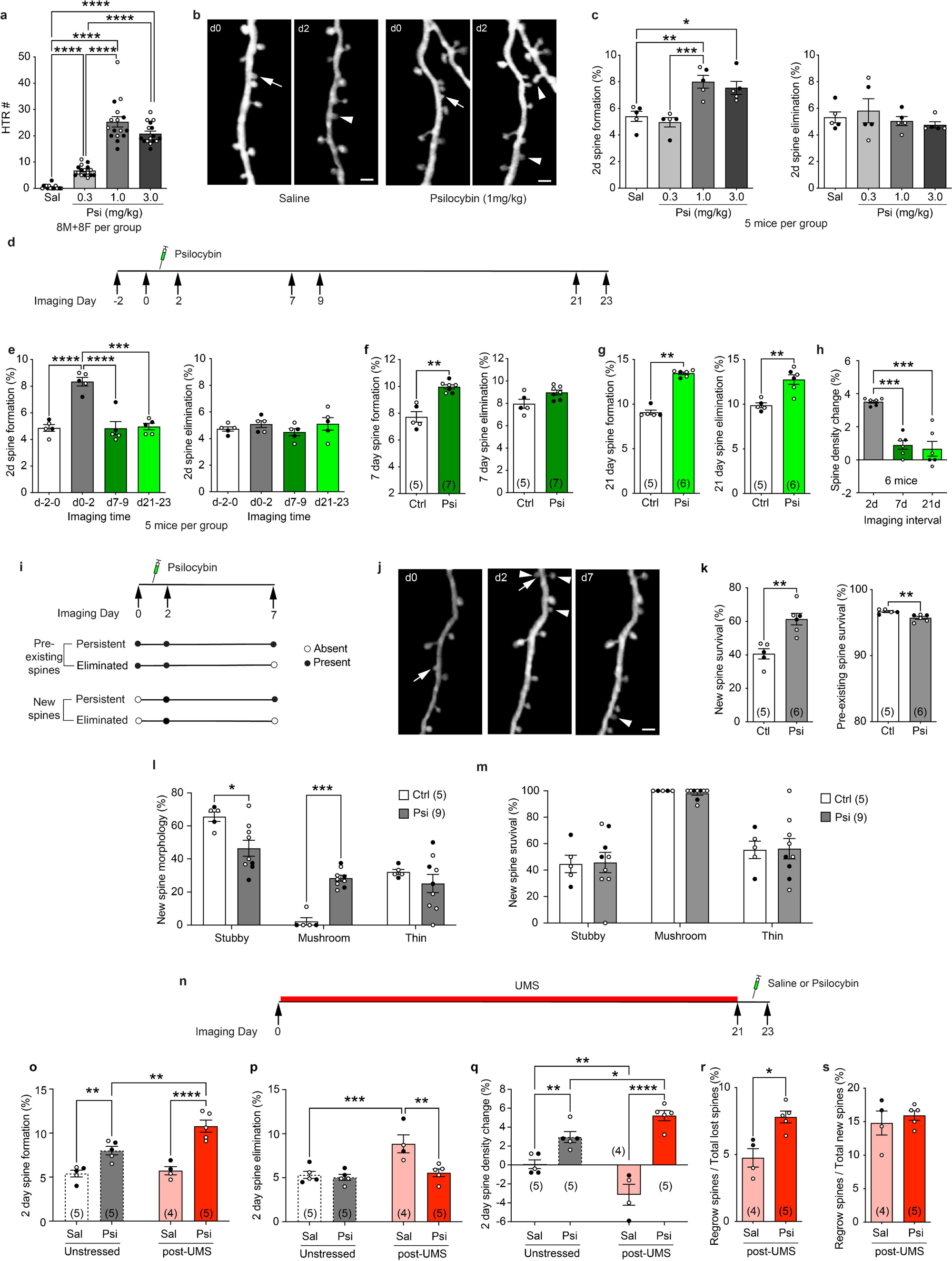
Psilocybin induces dose-dependent hallucinogenic responses and structural plasticity in mouse cortical circuits. **a**, HTRs following different doses of psilocybin treatment. Welch’s one-way ANOVA *W*(3,27.83) = 201.8, *p* < 0.0001, followed by Dunnett’s T3 multiple comparisons test. Sal: saline-treated; Psi: psilocybin-treated. **b**, Examples of *in vivo* 2P imaging of 2d spine dynamics with saline and psilocybin treatment. Arrows: spine elimination; arrowheads: spine formation. Scale bar: 2 µm. **c**, Structural dynamics of spines over 2d in response to different doses of psilocybin. Formation: one-way ANOVA *F*(3,16) = 12.43, *p* = 0.0002, followed by Tukey’s multiple comparisons test. Elimination: one-way ANOVA *F*(3,16) = 0.7065, *p* = 0.562. **d**, Timeline of longitudinal 2P imaging and psilocybin administration. **e**, 2d spine dynamics at different times before and after psilocybin treatment. Formation: repeated measures one-way ANOVA *F*(3,12) = 23.03, *p* < 0.0001, followed by Tukey’s multiple comparisons test. Elimination: repeated measures one-way ANOVA *F*(3,12) = 0.8061, *p* = 0.5144. **f**, 7d spine dynamics in control (Ctrl) vs Psi mice. Mann-Whitney test, formation: *U* = 0, *p* = 0.0025; elimination: *U* = 6, *p* = 0.0682. **g**, 21d spine dynamics in Ctrl vs Psi mice. Mann-Whitney test, formation *U* = 0, *p* = 0.0043; elimination *U* = 1, *p* = 0.0087. **h**, Spine density changes after psilocybin treatment over time. Repeated measures one-way ANOVA *F*(2,10) = 25.27, *p* = 0.0001, followed by Tukey’s multiple comparisons test. **i**, Schematic of the spine fate analysis. **j**, An example illustrating different fates of new spines. **k**, 7d survival rate in Ctrl vs Psi mice. Mann-Whitney test, new spines *U* = 0, *p* = 0.0043; pre-existing spines *U* = 0.5, *p* = 0.0065. **l**, Percentage of new spines in different morphological categories. Repeated measures two-way ANOVA, main effect of morphology *F*(2,24) = 30.26, *p* < 0.0001; main effect of treatment *F*(1, 12) = 0.5357, *p* = 0.4783; interaction *F*(2,24) = 9.65, *p* = 0.0008; followed by Šídák’s multiple comparisons test (Ctrl vs Psi). **m**, Survival rates of new spines in different morphological categories. Repeated measures two-way ANOVA, main effect of morphology *F*(2,24) = 24.15, *p* < 0.0001; main effect of treatment *F*(1,12) = 0.5357, *p* = 0.4783; interaction *F*(2,24) = 0.0219, *p* = 0.9784. **n**, Timeline of longitudinal 2P imaging with UMS and psilocybin administration. **o-q,** 2d spine formation (**o**), elimination (**p**), and spine density changes (**q**) in Sal vs Psi post-UMS mice in comparison with unstressed mice (same data as in Fig. 1c). Two-way ANOVA followed by uncorrected Fisher’s LSD test. Formation (**o**): main effect of Psi treatment *F*(1,15) = 52.72, *p* < 0.0001; main effect of UMS *F*(1,15) = 8.674, *p* = 0.01; interaction *F*(1,15) = 5.422, *p* = 0.0343. Elimination (**p**): main effect of Psi treatment *F*(1,15) = 10.28, *p* = 0.0059; main effect of UMS *F*(1,15) = 13.23, *p* = 0.0024; interaction *F*(1,15) = 7.307, *p* = 0.0164. Spine density change (**q**): main effect of Psi treatment *F*(1,15) = 70.19, *p* < 0.0001; main effect of UMS *F*(1,15) = 0.5218, *p* = 0.4812; interaction *F*(1,15) = 16.72, *p* = 0.0010. **r**, Percentage of lost spines that regrew in Sal vs Psi post-UMS mice. Mann-Whitney test, *U* = 0, *p* = 0.0159. **s**, Percentage of new spines that grew at sites of previous spine elimination in Sal vs Psi post-UMS mice. Mann-Whitney test, *U* = 9, *p* = 0.9127. Hereinafter *n* = number of mice, males are shown as filled circles and females as open circles. \**p* < 0.05, \*\**p* < 0.01, \*\*\**p* < 0.001, \*\*\*\**p* < 0.0001.

We also examined the neuroplastic effects of psilocybin through *in vivo* two-photon (2P) imaging of dendritic spines in *Thy1*-GFP-M mice, which express cytoplasmic green fluorescent protein predominantly in a sparse subset of cortical L5 PyrNs^25^ (Fig. 1b, Extended Data Fig. 2a). Mice were first imaged at baseline (d0), received either psilocybin (0.3, 1, or 3 mg/kg) or saline injection after 1 day (d1), and were re-imaged 24 h later (d2). Compared with saline controls, 1 and 3 mg/kg doses both increased spine formation significantly to a similar level, whereas the 0.3 mg/kg dose did not; spine elimination over the same period was unaffected across all doses (Fig. 1c). Consequently, 1 and 3 mg/kg doses led to a net gain in spine density over the 2d interval (Extended Data Fig. 2b). As both the HTR and neuroplastic effects of psilocybin plateaued at 1 mg/kg, we focused on this dose in all subsequent studies. In an independent cohort, the same mice were imaged during baseline and after 1 mg/kg psilocybin, which revealed no sex difference in baseline or psilocybin-induced spine dynamics (Extended Data Fig. 2c). A dendrite-level analysis further showed that both baseline and psilocybin-induced spine formation occurred broadly across imaged dendritic segments, rather than concentrating on a subset of highly plastic dendrites (Extended Data Fig. 3).

Next, we investigated the persistence of neuroplasticity induced by a single dose of psilocybin by following spine dynamics on the same dendritic segments before and after psilocybin treatment for more than three weeks (Fig. 1d). While spine formation between d0 and d2 significantly increased, neither spine formation nor elimination over a 2d interval at 1- or 3-week post-treatment differed significantly from the pre-treatment baseline (Fig. 1e), suggesting that psilocybin only transiently enhances spine formation. However, when measured over a 7d or a 21d interval, spine formation remained elevated in psilocybin-treated mice (Fig. 1f,g). Interestingly, spine elimination in psilocybin-treated mice significantly increased over 21d (Fig. 1g). Thus, the magnitude of net spine density changes diminished as the imaging interval increased (Fig. 1h). To assess how the new spines formed immediately after psilocybin treatment contribute to long-lasting alterations in synaptic circuits, we classified spines as either pre-existing (present on both d0 and d2 of imaging) or newly formed (present on d2 but not on d0) and tracked their fate on d7 and d21 (Fig. 1i, Extended Data Fig. 4a). In control mice, most new spines were unstable: only 40.6 ± 3.0% survived to d7 and 29.3 ± 4.1% to d21. After psilocybin treatment, however, a significantly higher fraction of new spines persisted over the same intervals (61.3 ± 3.5% to d7; 48.0 ± 3.4% to d21). On the contrary, pre-existing spines in psilocybin-treated mice were less likely to survive than their counterparts in control mice (Fig. 1j,k, Extended Data Fig. 4b). These results suggest that, although it only transiently boosts spine formation, psilocybin alters existing neural circuits with a long-lasting impact.

As spine morphology correlates with synaptic strength and dynamics^26,27^, we next investigated the morphological distribution of newly formed spines and their fate. In control mice, most new spines were stubby (65.6 ± 2.9%), and few were mushroom spines (2.2 ± 2.2%), with the remainder being thin spines (Fig. 1l). By d7, 55.4 ± 6.7% of stubby spines were lost, whereas almost all mushroom spines persisted (Fig. 1m). In psilocybin-treated mice, however, stubby spines accounted for a significantly smaller fraction of new spines (46.5 ± 4.8%), and mushroom spines a significantly larger fraction (28.4 ± 1.8%; Fig. 1l). The survival rates of stubby and mushroom spines were comparable to those in saline controls (Fig. 1m). Previous studies suggest that mushroom spines are more mature than stubby spines^26,28^. Thus, our data suggests that psilocybin not only promotes spine formation but also accelerates spine maturation.

To determine whether psilocybin-induced synaptic remodeling also occurs in the stressed brain, we subjected mice to 21d of unpredictable mild stress (UMS) and administered either saline or psilocybin (1 mg/kg) 1d after termination of stress. Dendritic spines were imaged before UMS, at the end of UMS, and again 1d after saline or psilocybin administration (Fig. 1n). We found that psilocybin drove a greater increase in spine formation in post-UMS mice compared to unstressed controls (Fig. 1o). Furthermore, while post-UMS mice continued to exhibit elevated spine elimination even after stress cessation, psilocybin reduced this spine elimination to levels comparable to those in unstressed controls (Fig. 1p). Consequently, psilocybin induced a greater net gain in spine density in post-UMS mice than in unstressed mice (Fig. 1q). Together, these data suggest that post-stress psilocybin treatment not only promotes new spine formation but also stabilizes existing spines.

Are new spines induced by post-UMS psilocybin treatment preferentially formed at sites of previous spine elimination during UMS? In psilocybin-treated mice, a significantly higher fraction of spines lost during UMS were recovered, compared to saline-treated post-UMS mice (7.8 ± 0.4 % vs. 4.8 ± 0.7%, *p* = 0.0159, Fig. 1r). However, among new spines, the percentage that emerged at sites of previous spine elimination was comparable between psilocybin- and saline-treated groups (psilocybin 15.9 ± 0.7 % vs. saline 14.8 ± 1.8 %, *p* = 0.9127, Fig. 1s). Thus, the enhanced recovery of lost spines following psilocybin treatment is likely driven by an overall increase in spine formation, rather than the selective targeting of sites of previous spine elimination. Furthermore, in psilocybin-treated post-UMS mice, a significantly higher proportion of new spines assumed the mushroom morphology, compared to saline-treated post-UMS mice (Extended Data Fig. 5), suggesting that psilocybin also induces rapid structural maturation of new spines in the post-stress brain.

We next investigated how 5-HT_2A_R signaling in L5 PyrNs contributes to psilocybin’s hallucinogenic and neuroplastic effects, by either restricting 5-HT_2A_R expression exclusively to these neurons or abolishing 5-HT_2A_R expression specifically in these neurons. First, we used a 5-HT_2A_R knock-out conditional rescue line (*htr2a*^stop/stop^), in which a “stop” cassette flanked by *lox-P* sites was inserted between the promoter and the first exon of the *htr2a* gene to prevent its transcription^29^. When crossed with Rbp4-Cre mice, which express Cre recombinase in L5 PyrNs of both pyramidal tract (PT) and intratelencephalic (IT) subtypes across cortical areas^30–32^, 5-HT_2A_R expression is restored in Cre+ neurons. Hereafter we refer to *htr2a*^stop/stop^ mice as 5-HT_2A_R full knock-out (FKO) mice, and Rbp4-Cre^+^; *htr2a*^stop/stop^ mice as conditional rescue (CR) mice. Immunohistochemistry confirmed the abolition and restoration of 5-HT_2A_R expression in FKO and CR mice, respectively (Fig. 2a). As expected, psilocybin failed to elicit the HTR in FKO mice (Fig. 2b). Interestingly, it failed to elicit the HTR in CR mice as well (Fig. 2b), indicating that 5-HT_2A_R signaling in L5 PyrNs alone is insufficient to induce this hallucination-like behavior.

**Figure 2.**
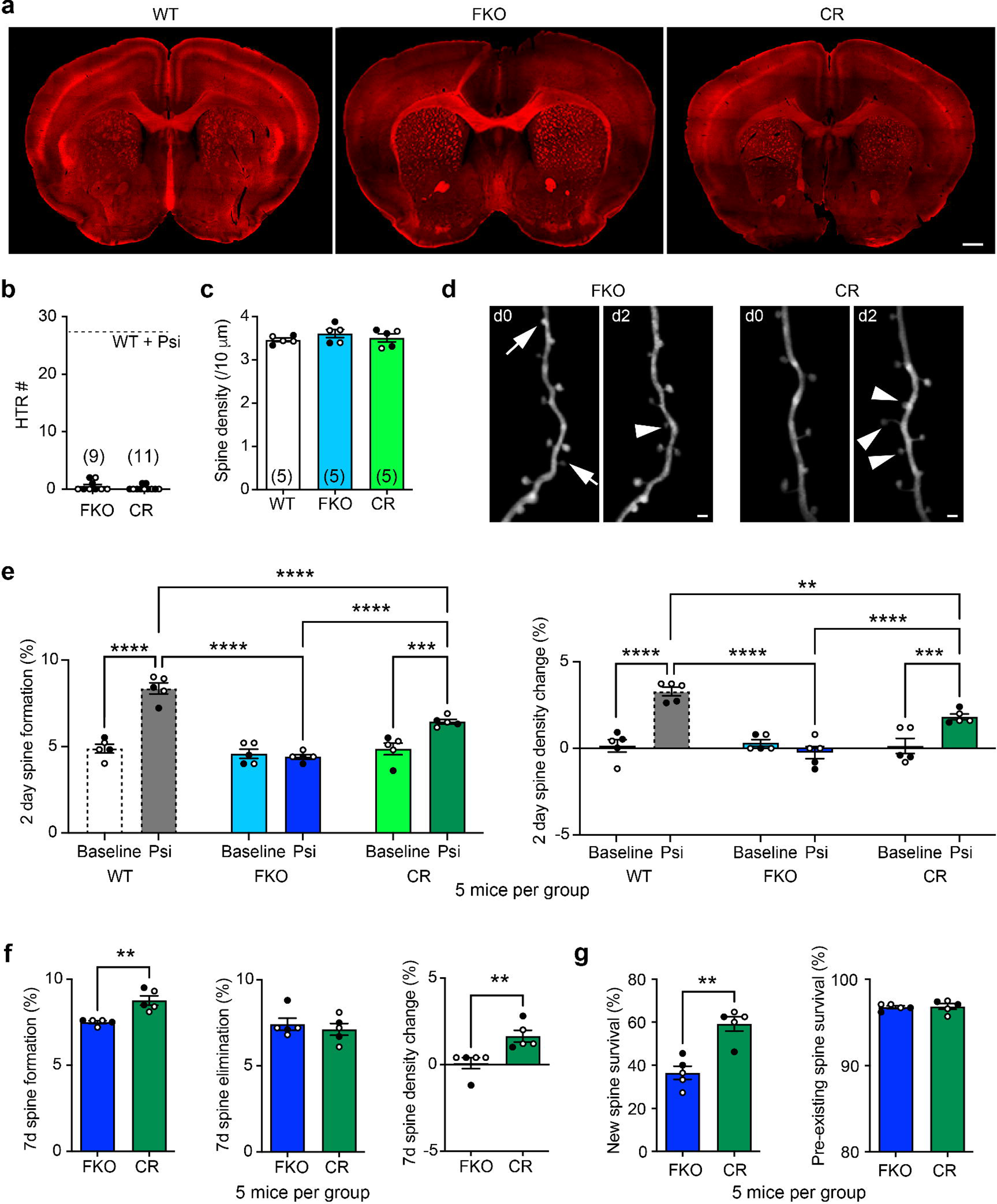
Selective expression of 5-HT_2A_Rs in cortical L5 PyrNs suffices to restore psilocybin-induced spine formation and stabilization but not HTRs. **a**, Immunohistochemistry showing the abolition and restoration of 5-HT_2A_R expression in L5 PyrNs in FKO and CR mice, respectively. Scale bar: 500 µm. **b**, 1 mg/kg psilocybin does not elicit HTRs in FKO or CR mice. **c**, Spine density on the apical dendritic tufts of L5 PyrNs in WT, FKO, and CR mice. One-way ANOVA *F*(2,12) = 0.8753, *p* = 0.4417. **d**, Example of *in vivo* 2P imaging of 2d spine dynamics in FKO and CR mice with 1 mg/kg psilocybin treatment. Arrows: spine elimination; arrowheads: spine formation. Scale bar: 2 µm. **e**, 2d spine formation and density changes in WT, FKO, and CR mice under baseline and with psilocybin treatment. Two-way ANOVA followed by uncorrected Fisher’s LSD test. Formation: main effect of treatment *F*(1,24) = 61.73, main effect of genotype *F*(2,24) = 34.73, interaction *F*(2,24) = 26.05, *p* < 0.0001 for all. Spine density changes: main effect of treatment *F*(1,24) = 31.73, main effect of genotype *F*(2,24) = 14.37, interaction *F*(2,24) = 18.22, *p* < 0.0001 for all. **f**, 7d spine dynamics in FKO vs CR mice with psilocybin treatment. Mann-Whitney test, formation *U* = 0, *p* = 0.0079; elimination *U* = 11, *p* = 0.8333; spine density changes *U* = 0, *p* = 0.0079. **g**, 7d survival rate of spines in FKO vs CR mice with psilocybin treatment. Mann-Whitney test, new spines *U* = 0, *p* = 0.0079; pre-existing spines *U* = 10, *p* = 0.6349.

The spine density in both FKO and CR mice was comparable to that in WT mice (Fig. 2c), suggesting no gross deficit in synaptic development. In FKO mice, psilocybin did not alter spine formation or elimination over 2d. In contrast, psilocybin significantly increased spine formation in CR mice over 2d, albeit to a lesser extent than in psilocybin-treated WT mice; spine elimination was unaffected, leading to a net gain in spine density (Fig. 2d,e, Extended Data Fig. 6a). Similarly, psilocybin treatment resulted in higher spine formation and spine density gain over 7d in CR mice than in FKO mice (Fig. 2f). New spines formed in CR mice were more likely to persist to d7 than those in FKO mice (Fig. 2g), with a survival rate comparable to psilocybin-induced new spines in WT mice (Extended Data Fig. 6b). Interestingly, after psilocybin treatment, the survival rate of pre-existing spines in CR mice was comparable to that in FKO mice (Fig. 2g) and higher than that in WT mice (Extended Data Fig. 6c). These findings suggest that the selective expression of 5-HT_2A_R in L5 PyrNs suffices to promote spine formation and stabilization without affecting pre-existing synaptic connections.

Is 5-HT_2A_R signaling in cortical L5 PyrNs necessary for either psychedelic-induced HTRs or neuroplasticity? To address this question, we selectively abolished 5-HT_2A_R expression in L5 PyrNs by crossing a 5-HT_2A_R conditional knock-out line (*htr2a*^flox/flox^)^33^ with Rbp4-Cre mice to generate Rbp4-Cre^+^; *htr2a*^flox/flox^ mice (hereafter referred to as CKO mice). The *htr2a*^flox/flox^ mice responded to psilocybin with HTRs as expected (Fig. 3a). Surprisingly, CKO mice also exhibited HTRs upon psilocybin treatment (Fig. 3a), indicating that 5-HT_2A_R signaling in cortical L5 PyrNs is not necessary for the HTR. Spine densities in *htr2a*^flox/flox^ and CKO mice were normal (compare Fig. 3b with Fig. 2c). In CKO mice psilocybin altered neither spine formation nor elimination, while in *htr2a*^flox/flox^ mice it elevated spine formation and spine density over 2d as expected (Fig. 3c,d). Similarly, psilocybin treatment resulted in higher spine formation and net spine density gain over 7d in *htr2a*^flox/flox^ mice, compared to CKO mice (Fig. 3e). In *htr2a*^flox/flox^ mice, new spines were also more likely to persist to d7 than in CKO mice, but pre-existing spines had the opposite fate (Fig. 3f). Together with the findings above, these results suggest that 5-HT_2A_R signaling in L5 PyrNs is both necessary and sufficient for psilocybin’s neuroplastic effects (promoting synapse formation and stabilization), but neither necessary nor sufficient for psilocybin-induced HTRs.

**Figure 3.**
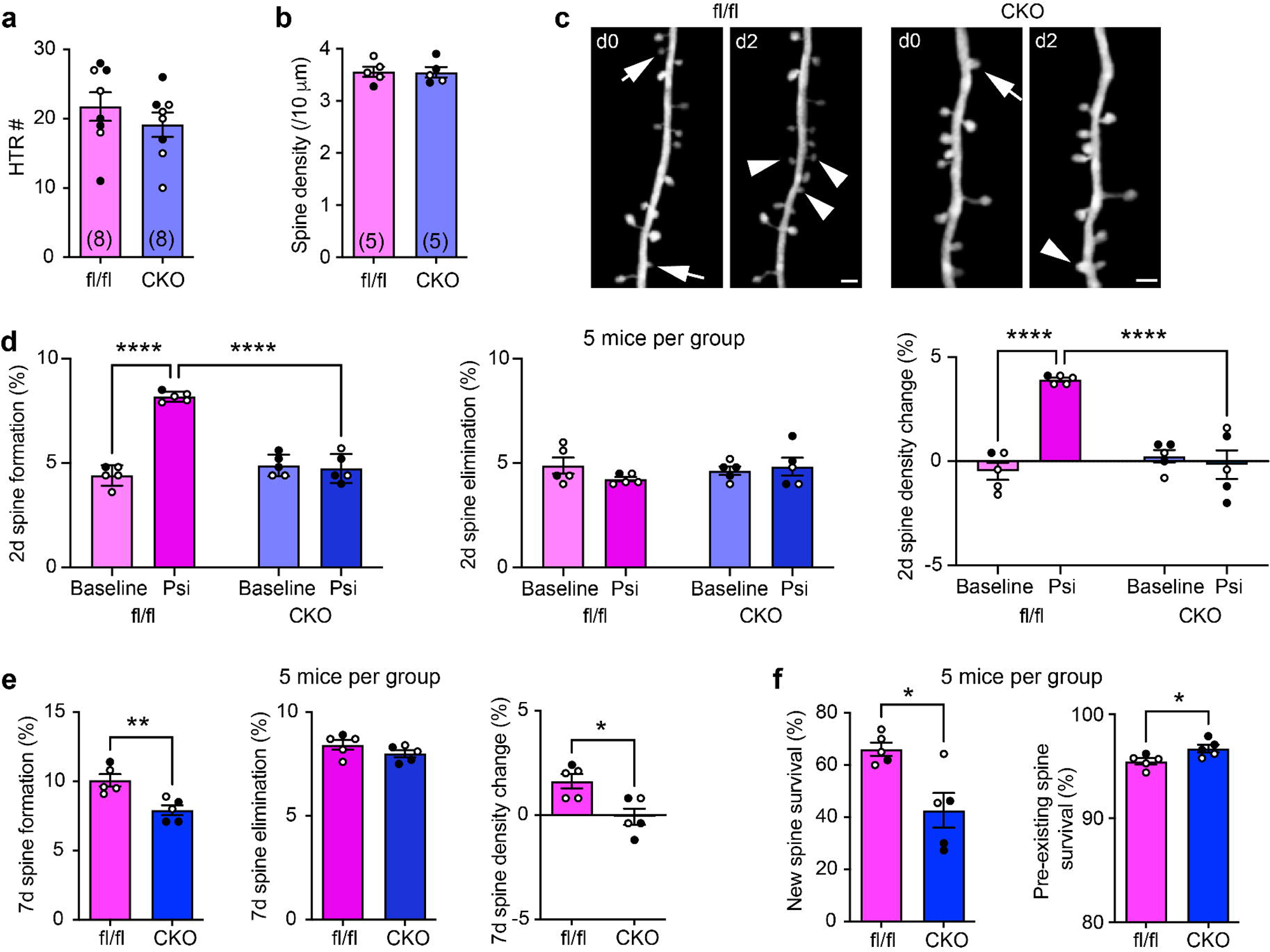
Selective knock-out of 5-HT_2A_Rs in cortical L5 PyrNs prevents psilocybin-induced spine formation but not HTRs. **a**, 1 mg/kg psilocybin elicits HTRs in *htr2a*^flox/flox^ (fl/fl) and CKO mice comparably. Mann-Whitney test, *U* = 22.50, *p* = 0.3414. **b**, Spine density on the apical dendritic tufts of L5 PyrNs in fl/fl and CKO mice. Mann-Whitney test, *U* = 12, *p* > 0.9999. **c**, Example of *in vivo* 2P imaging of 2d spine dynamics in fl/fl and CKO mice with 1 mg/kg psilocybin treatment. Arrows: spine elimination; arrowheads: spine formation. Scale bar: 2 µm. **d**, 2d spine dynamics in fl/fl vs CKO mice under baseline and with psilocybin treatment. Two-way ANOVA followed by uncorrected Fisher’s LSD test. Formation: main effect of treatment *F*(1, 16) = 63.27, main effect of genotype *F*(1,16) = 41.84, interaction *F*(1,16) = 73.38, *p* < 0.0001 for all. Elimination: main effect of treatment *F*(1,16) = 0.5039, *p* = 0.488; main effect of genotype *F*(1,16) = 0.3373, *p* = 0.5695; interaction *F*(1,16) = 1.837, *p* = 0.1942. Spine density changes: main effect of treatment *F*(1,16) = 21.68, *p* = 0.0003; main effect of genotype *F*(1,16) = 15.30, *p* = 0.0012; interaction *F*(1, 16) = 31.22, *p* < 0.0001. **e**, 7d spine dynamics in fl/fl vs CKO mice with psilocybin treatment. Mann-Whitney test, formation *U* = 0, *p* = 0.0079; elimination *U* = 6, *p* = 0.1984; spine density changes *U* = 2, *p* = 0.0476. **f**, 7d survival rate of spines in fl/fl vs CKO mice with psilocybin treatment. Mann-Whitney test, new spines *U* = 2, *p* = 0.0317; pre-existing spines *U* = 2, *p* = 0.0397.

Our study revealed several features of psilocybin-induced structural dynamics in synaptic circuits. The elevation in spine formation is rapid but transient, consistent with previous reports^13,17^. Notably, mushroom-shaped spines account for a significantly larger fraction of new spines in psilocybin-treated mice than in controls. Characterized by large spine heads and expanded postsynaptic densities, which correlate strongly with AMPA receptor content and synaptic strength^34,35^, mushroom spines are believed to host mature and stable synapses^26,28^. Thus, psilocybin not only promotes synapse formation but also accelerates synapse maturation. Contrary to a previous report^13^, we found that psilocybin-induced new spines are preferentially stabilized. This discrepancy may arise from regional differences in spine dynamics, as we imaged the somatosensory cortex, whereas the other study focused on the frontal cortex (Cg1/M2). Moreover, we observed a concomitant elevation in the elimination of pre-existing spines, which leads to a cumulative increase in spine elimination that gradually drives spine density towards the pre-treatment level in the normal brain.

In the post-stress brain, psilocybin exerts a more pronounced structural effect than in the normal brain, inducing a greater amount of spine formation and normalizing the elevated spine elimination that persists after stress cessation, leading to a larger net gain in spine density. Importantly, although psilocybin promotes the recovery of spines lost during stress, new spines do not preferentially emerge at sites of previous spine elimination. This pattern differs from the reported effects of ketamine on corticosterone-treated mice^36^. Whereas the prevalent view emphasizes the importance of a sustained increase in spine density as a physical substrate for the durable therapeutic effect of psilocybin and ketamine for depression^5^, our findings suggest a more nuanced view that psilocybin induces global synaptic circuit remodeling in a manner that depends on the context (*e.g.*, normal or pathological) in which psilocybin is administered.

Our findings further support a cellular dissociation between psilocybin-induced behavioral and neuroplastic effects, consistent with a recent report that 5-HT_2A_R expression in glutamatergic neurons in the medial prefrontal cortex is dispensable for the psilocybin-induced HTR but required for the structural plasticity of PT-type L5 PyrNs^17^. The Rbp4-Cre line used in our study expresses Cre recombinase in both PT and IT subtypes of L5 PyrNs^30–32^, but only labels about 50% of all L5 PyrNs^37,38^. When *Thy1*-GFP-M line is crossed with Rbp4-Cre line, almost all GFP+ L5 PyrNs are Cre+, as revealed by immunohistochemistry (Extended Data Fig. 7). We expect that most of the GFP+ neurons we imaged are PT, as supported by a connectomic study^39^ that mapped the whole-brain axonal projections of individual neurons in *Thy1*-GFP-M mice, and by a recent transcriptomic study^40^ suggesting that about two-thirds of Rbp4-Cre+ neurons are PT.

Recent studies have reported inconsistent results on the necessity of 5-HT_2A_R signaling in psilocybin-induced neuroplasticity^13–17^. Using cell type-specific knock-out and rescue strategies, we demonstrated that 5-HT_2A_R signaling in cortical L5 PyrNs is both necessary and sufficient for their synaptic remodeling in response to psilocybin. Selective restoration of 5-HT_2A_R expression in L5 PyrNs (CR mice) partially rescues psilocybin-induced increases in spine formation and stabilization, whereas selective deletion of 5-HT_2A_Rs from these neurons (CKO mice) abolishes these effects. Intriguingly, while psilocybin accelerates the elimination of pre-existing spines in WT mice, it fails to do so in both CR and CKO mice. These results suggest that 5-HT_2A_R signaling in L5 PyrNs accounts for a large component of psilocybin-induced spine formation and stabilization, although non-cell-autonomous contributions from other types of cells that normally also express 5-HT_2A_Rs cannot be neglected. The reorganization of pre-existing neural circuits, on the other hand, results from the synergy between elevated spine formation and the action of psilocybin on other cell types.

Moreover, CR mice fail to exhibit HTRs following psilocybin treatment, in contrast to a previous report that selective restoration of 5-HT_2A_R expression in cortical excitatory neurons suffices to rescue psychedelic-induced HTRs^41^. In addition, CKO mice still exhibit psilocybin-induced HTRs. These results suggest that although L5 PyrNs are the predominant type of cortical neuron expressing 5-HT_2A_Rs, the HTR likely involves additional cell types and brain circuits. We note that, although the HTR is widely used as a behavioral proxy for the hallucinogenic potential of serotonergic psychedelics, this stereotyped motor output does not measure perceptual experiences; thus, our results should not be interpreted as directly addressing the neural basis of psychedelic-induced hallucinations in humans.

Nevertheless, recent developments in medicinal chemistry corroborate the idea that 5-HT_2A_R-related neuroplasticity can be separated from hallucination-like behavioral manifestations. Tabernanthalog (TBG), a non-hallucinogenic structural analog of 5-methoxy-*N,N*-dimethyltryptamine, promotes cortical dendritic spine formation at baseline and in mouse models of stress^18,42,43^; such neuroplastic effects are mediated through biochemical pathways involving 5-HT_2A_R, TrkB, mTOR and AMPA receptor activation as classical hallucinogenic psychedelics^42^. Similarly, the lysergic acid diethylamide (LSD) analog 2-Br-LSD, a partial agonist of 5-HT_2A_Rs, does not induce the HTR in mice but promotes dendritogenesis and spinogenesis in cultured cortical neurons^22^. Additional compounds, including (+)-JRT, an LSD analog with reduced hallucinogenic potential, and zalsupindole (also known as AAZ-A-154 or DLX-001), a non-hallucinogenic iso-tryptamine psychoplastogen, have also been reported to promote cortical spinogenesis and neuritogenesis while minimizing psychedelic-like behavioral effects^19,44,45^. Taken together, our findings indicate that psilocybin’s neuroplastic and hallucinogenic effects are dissociable at the cellular level, which informs the potential to develop neuroplasticity-based therapies without undesirable psychotropic effects.

## Methods

### Experimental animals

C57BL/6J (JAX# 000664) and *Thy1*-GFP-M (JAX# 007788) mice were originally purchased from The Jackson Laboratory; the Rbp4-Cre (MMRRC 031125-UCD) mouse line was obtained from Dr. Lu Chen (Stanford University); the 5-HT_2A_R knock-out conditional rescue mouse line (*htr2a*^stop/stop^)^29^ was obtained from Dr. David E. Olson (University of California, Davis); the 5-HT_2A_R conditional knock-out mouse line (*htr2a*^flox/flox^)^33^ was obtained from Dr. Hail Kim (Korea Advanced Institute of Science and Technology). To generate conditional rescue (CR) and conditional knockout (CKO) mice for *in vivo* imaging, we used the following breeding strategy. To generate CR mice, homozygous htr2a^stop/stop^ mice were first crossed separately with *Thy1*-GFP-M mice or with Rbp4-Cre+ mice. The resulting offspring were crossed back with homozygous htr2a^stop/stop^ mice to generate htr2a^stop/stop^;GFP+ and Rbp4-Cre+;htr2a^stop/stop^ mice, respectively. These two lines were then crossed to obtain Rbp4-Cre+;htr2a^stop/stop^;GFP+ (*i.e.*, CR) and Rbp4-Cre-;htr2a^stop/stop^;GFP+ (*i.e.*, full knockout or FKO) offspring. To generate CKO mice, an analogous breeding scheme was adopted for homozygous htr2a^flox/flox^ mice, to generate Rbp4-Cre+;htr2a^flox/flox^;GFP+ (*i.e.*, CKO) and Rbp4-Cre-;htr2a^flox/flox^;GFP+ (*i.e.*, floxed control) mice. Mice are group-housed on a 12 h light/dark cycle and randomly assigned to experimental groups. All animal experiments were carried out in accordance with protocols approved by the Institutional Animal Care and Use Committee of University of California, Santa Cruz.

### Drug preparation and administration

Psilocybin was administered through intraperitoneal injection at a dosage of 0.3, 1, or 3 mg/kg of bodyweight. USP-grade saline (0.9%) was used as the vehicle.

### Head-twitch response (HTR) recording and annotation

The mouse was placed in an empty standard mouse cage after psilocybin or vehicle injection. Its behavior was recorded with an iPhone 15 equipped with a 48-MP main sensor camera (fps = 30) for 20 min. Behavioral videos were manually annotated for the HTR using the BORIS software^46^. The annotator was blind to the animal’s experimental condition.

### Cranial window implantation

We performed cranial window implantation on mice around P60 according to established protocols^47^ with slight modifications. In brief, the mouse was anesthetized with isoflurane (4% for induction, 1.5% for maintenance). Ophthalmic ointment was applied to prevent eye desiccation and irritation; dexamethasone (2 mg/kg bodyweight) was injected into the quadriceps, and carprofen (5 mg/kg bodyweight) was injected intraperitoneally. A circular piece of the skull over the somatosensory cortex (centered approximately at AP −1.5 mm, ML 1.5 mm) was removed with a trephine (Fine Science Tools, diameter = 2.3 mm) driven by a high-speed micro-drill (Foredom K1070). The cranial window was sealed with an imaging port made of a round glass coverslip (#2, diameter = 2.3 mm) glued to an overlaying annular glass “doughnut” (#1, inner diameter = 2 mm, outer diameter = 3 mm, Potomac Photonics, Inc.). Dental cement (Jet Denture Repair, Lang Dental) was applied over the exposed skull to secure a custom-made stainless-steel head plate onto the skull. The mouse received the antibiotic enrofloxacin (5 mg/kg) and the analgesic buprenorphine (0.3 mg/kg) preemptively and then daily for 2 more days.

### *In vivo* two-photon (2P) imaging of dendritic spines and data analysis

*In vivo* 2P imaging of dendritic spines was performed on a 2P microscope (Ultima IV, Bruker Co.) equipped with a 40x/NA = 0.8 water immersion objective (Olympus) and an ultrafast 2P laser (Mai Tai HP, Spectra-Physics) operating at 940 nm. The mouse was anaesthetized with an intraperitoneal injection of a mixture of 17 mg/ml ketamine and 1.7 mg/ml xylazine in 0.9% saline (5.0 ml/kg) and mounted on a custom-made stage for imaging. Stacks of images were acquired with a Z-step size of 1 µm at 4x digital zoom. Relocation of the same dendrites in subsequent imaging sessions was achieved by reference to blood vessels and the dendritic branching pattern. Data analysis was performed on 3D image stacks in ImageJ as described previously^48^. In *Thy1*-GFP-M line, L2/3 neurons are occasionally labeled with GFP, and some of their dendritic segments may extend into cortical L1. We therefore selected imaging regions in which no GFP+ L2/3 neuronal somata were visible at the imaging site and focused on dendritic segments consistent with the apical tuft dendrites of L5 PyrNs within the first 150 μm below the pial surface (which primarily corresponds to cortical L1). We acknowledge that this approach cannot exclude every possible L2/3 neuron contribution with absolute certainty, but the scarcity of L2/3 neurons labeled in this line (see Extended Data Fig. 2a) together with our selection criterion ensure that the imaged dendritic segments are predominantly the apical tufts of L5 PyrNs. Typically, 200-250 spines were analyzed per animal per session. The number of dendritic segments and initial spine counts of each mouse in each figure are given in Supplementary Table 1. The percentage of spines formed/eliminated was calculated as the number of spines formed/eliminated divided by the total number of spines counted from the previous imaging session.

Morphological categorization of spines was performed according to criteria described previously^49^. Specifically, the analyst scrolled through the z-stack to evaluate the presence of a spine neck and the relative size of the spine head. Mushroom spines were defined as protrusions with a spine head width greater than twice the neck width. Stubby spines were defined as short protrusions less than 0.5 μm in length without a discernible neck. Thin spines were defined as protrusions longer than 0.5 μm with a head width less than or equal to twice the neck width. When a spine protruded above or below the dendritic shaft, we classified it as a mushroom spine only if it appeared narrower/dimmer in at least one z-plane between the dendritic shaft and the bulbous spine head, reminiscent of a neck-like structure. Otherwise, it would be classified as a stubby spine (thin spines protruding above or below the dendritic shaft would be hardly visible). We acknowledge that the inferior axial resolution makes morphological classification in this case less reliable, leading to a potential underestimation of mushroom and thin spines. Nevertheless, such cases are relatively few, so we do not expect them to impact the statistical analysis significantly.

We also investigated whether psilocybin-induced spine formation was uniformly distributed across dendritic segments or concentrated on a small subset of dendrites, using the following permutation test. Let *D* denote the total number of dendritic segments, *n_i_* the initial spine count of the *i*^th^ dendrite, and 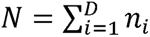 the total initial spine count. Let *k_i_* denote the number of new spines formed on the *i*^th^ dendrite, yielding a total of 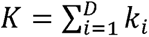 new spines across all dendrites. Under the null hypothesis of uniform spine formation, the probability of a new spine being formed on the *i*^th^ dendrite is proportional to the initial spine count on the dendrite, which gives the null probability distribution *Q*, where *Q_i_* = *n_i_*/*N*. In each simulated trial, we randomly and independently distributed exactly *K* new spines among the *D* dendrites according to the probability distribution *Q*. For each trial, we calculated the simulated spine formation fraction for the *i*^th^ dendrite as 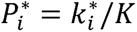, where 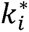 denotes the number of new spines assigned to this dendrite. Note that because 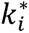 must be an integer, 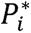 in general does not equal *Q_i_*. We quantified this deviation from uniformity using the Kullback-Leibler (KL) divergence between the simulated distribution *P*\* and the null distribution *Q* for each trial:

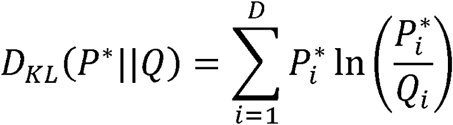

We also computed the KL divergence between the observed empirical distribution *P* (where *P_i_* = *k_i_*/*K*) and the null distribution *Q*. By comparing the empirical KL divergence given by *P* against the distribution of KL divergences generated from simulation (10,000 trials, each given by a *P*\*), we obtained a *p*-value. A significant *p*-value (*p* < 0.05) indicates that spine formation is significantly more concentrated on a subset of dendrites than would be expected by chance.

### Unpredictable mild stress (UMS)

Mice were subjected to UMS for 3 weeks with random stressors presented on a weekly rotating basis, as described previously^43^.

### Immunohistochemistry and fluorescence microscopy

The mouse was transcardially perfused with 4% paraformaldehyde (PFA) in 0.01M phosphate-buffered saline (PBS) as previously described^49^. The brain was post-fixed in 4% PFA at 4 overnight. For immunohistochemical detection of 5-HT_2A_R, the brain was cryoprotected in 30% sucrose solution, then sectioned into 40 µm thick coronal slices with a vibratome (Leica VT1000S). Free-floating slices were rinsed in PBS (10 min x 3), underwent antigen retrieval in 0.01M citrate buffer (sodium citrate 3 mg/ml, citric acid 0.4 mg/ml, pH = 6.0 [adjusted by NaOH]) at 95 for 5 min, rinsed again in PBS (10 min x 3), and immersed in the blocking solution (5% normal goat serum [NGS], 5% bovine serum albumin, 0.3% Triton X-100 in PBS) at room temperature for 2 h. Brain slices were then stained with a primary antibody against 5-HT_2A_R (rabbit polyclonal IgG fraction, Immunostar 24288, 1:320) in a solution of 5% NGS and 0.3% Triton X-100 in PBS at 4 for 48 h. They were rinsed in PBS (10 min x 3), stained with a biotinylated goat anti-rabbit secondary antibody (Vector Laboratories BA-1000-1.5, 1:1000) in a solution of 5% NGS in PBS at room temperature for 2 h, rinsed again in PBS (10 min x 3), stained with streptavidin conjugated with Alexa 594 (Invitrogen S32356, 1:1000) in a solution of 5% NGS in PBS at room temperature for 2 h, and finally rinsed in PBS (10 min x 3). Slices were counterstained with 4’-6-diamidino-2-phenylindole (DAPI; 1:36,000) for 10 min, rinsed in PBS (10 min x 3), and mounted on slides with Vectashield HardSet antifade mounting medium (Vector Laboratories H-1400-10). Brain slices were imaged on a Zeiss AxioImager Z2 widefield fluorescence microscope with a 10x/NA=0.45 air objective.

For immunohistochemical detection of Cre recombinase, the post-fixed brain was directly transferred into 1x PBS and sectioned into 40 µm thick coronal slices with a vibratome (Leica VT1000S). Free-floating slices were blocked in PBS containing 5% NGS and 0.5% Triton X-100 at room temperature for 1 h. Slices were then stained with a primary antibody against Cre recombinase (rabbit polyclonal, Synaptic Systems # 257 003, 1:1000) in a solution of 5% NGS in PBS at 4 °C overnight. After rinsing in PBS, slices were stained with an Alexa Fluor 594-conjugated goat anti-rabbit secondary antibody (Invitrogen A-11012, 1:500) at room temperature for 2 h, then counterstained with DAPI (1:36,000) for 10 min, rinsed in PBS (10 min x 3), and mounted on slides with Vectashield HardSet antifade mounting medium (Vector Laboratories H-1400-10). Brain slices were imaged on a Zeiss AxioImager Z2 widefield fluorescence microscope with ApoTome optical sectioning using a 10x/NA=0.45 air objective.

### Statistical analysis

All behavioral and spine dynamics were analyzed with the analyst blinded to the experimental conditions. All statistical analyses were performed using GraphPad Prism 10. Data from each mouse was treated as a single data point in a group. We report sample sizes, the statistical tests used, and the *p* values in figure, figure legends and Statistical Table. Statistical significance is defined as *p* < 0.05.

## Acknowledgements

We thank Dr. Aria Walls and Dr. Hyo Gun Lee for critical comments on this manuscript, Dr. Zi Ye for consultation on statistics, Stefan Abero for help with immunohistochemistry, and Dr. Benjamin Abrams (UCSC Life Sciences Microscopy Center) for technical support. This work was supported by grants from the National Institutes of Mental Health (R01MH127737 and R01MH136381) and National Institute on Aging (R01AG071787) to Y.Z.

## Author Contributions

J.L. and Y.Z. designed the study; J.J.B., E.K., S.M., and J.L. performed the experiments and data analysis; J.L. and Y.Z. wrote the manuscript.

## Data Availability

The data supporting the findings of this study are available from the corresponding authors upon reasonable request.

## Code Availability

The custom-written data analysis codes are available from the corresponding authors upon reasonable request.

