## Extended Data for "Dissociating the Hallucinogenic and Neuroplastic Effects of Psilocybin"

**Characterization of GFP+ cell distribution in the imaging region of *Thy1*-GFP-M mice**

In *Thy1*-GFP-M line, L2/3 neurons are occasionally labeled with GFP, and some of their dendritic segments may extend into cortical L1, where we performed *in vivo* 2P imaging. To further characterize the laminar distribution of GFP+ neurons in this line, we took 10 coronal sections (40 µm thick) of 3 mouse brains around the same location along the anterior-posterior axis as in *in vivo* spine imaging experiments. We examined a 500 µm wide cortical region in each slice, and in total found 459 GFP+ neurons with somata in L5, but only 14 GFP+ neurons with somata in L2/3. The distance of L2/3 neuronal somata from the pial surface ranged from 137 to 217 µm (average 177.6 ± 6.9 µm). An example illustrating the scarcity of GFP+ L2/3 neurons in contrast to L5 neurons is presented in Extended Data Fig. 2a.

**Dendrite-specific distribution of spine formation**

Previous studies have shown that spine remodeling can be spatially organized along dendrites, with learning-induced new spines forming in clusters and, in some cases, preferentially on selected dendritic branches^1,2^. We therefore analyzed spine formation at the level of individual dendritic segments to determine whether psilocybin-induced new spines were uniformly distributed across dendrites or concentrated on a small subset of dendrites. To avoid bias from very short dendritic segments, this analysis included only segments longer than 10 µm.

We first examined 2d spine formation in WT mice with and without 1 mg/kg psilocybin treatment. We tracked 186 dendritic segments from 9 psilocybin-treated mice (4M, 5F) and 188 segments from **9** untreated mice (4M, 5F). In both groups, a substantial fraction of dendritic segments exhibited no new spine formation; the remaining ones showed a wide range of apparent spine formation rates (from 3.8% **to 33.3%)**. It is noteworthy that even under the null hypothesis of uniform spine formation, these results are not surprising, as spine formation is a relatively sparse event (on average 8.1% and 5.1% for psilocybin-treated and untreated mice, respectively). Furthermore, the number of new spines formed on each dendritic segment was positively correlated with the initial spine count on that dendrite (Pearson *r* = 0.4134 for treated mice, *r* = 0.3504 for untreated mice, both *p* < 0.0001), consistent with globally distributed spine formation.

We next asked whether the observed spine formation was more concentrated on a subset of dendrites than expected by a uniform distribution. To address this question, we performed a permutation test to compare the empirical data with simulated data in which the same number of new spines were randomly and independently assigned to individual dendrites, with a probability proportional to the initial spine count on each dendrite. We used the Kullback–Leibler (KL) divergence to quantify how much the empirical or simulated new spine distribution deviated from the uniform distribution. We repeated this simulation 10,000 times to generate a reference distribution of KL divergence values and then compared the empirical KL divergence with this reference distribution to obtain a *p*-value. Using this approach, we analyzed 2d spine formation with and without 1 mg/kg psilocybin treatment in WT mice of both sexes. We found that the empirical KL divergence did not differ significantly from simulated data in all groups (Extended Data Fig. 3), supporting the idea that the observed spine formation is not driven by a small number of highly plastic dendrites.

**Extended Data Figures**

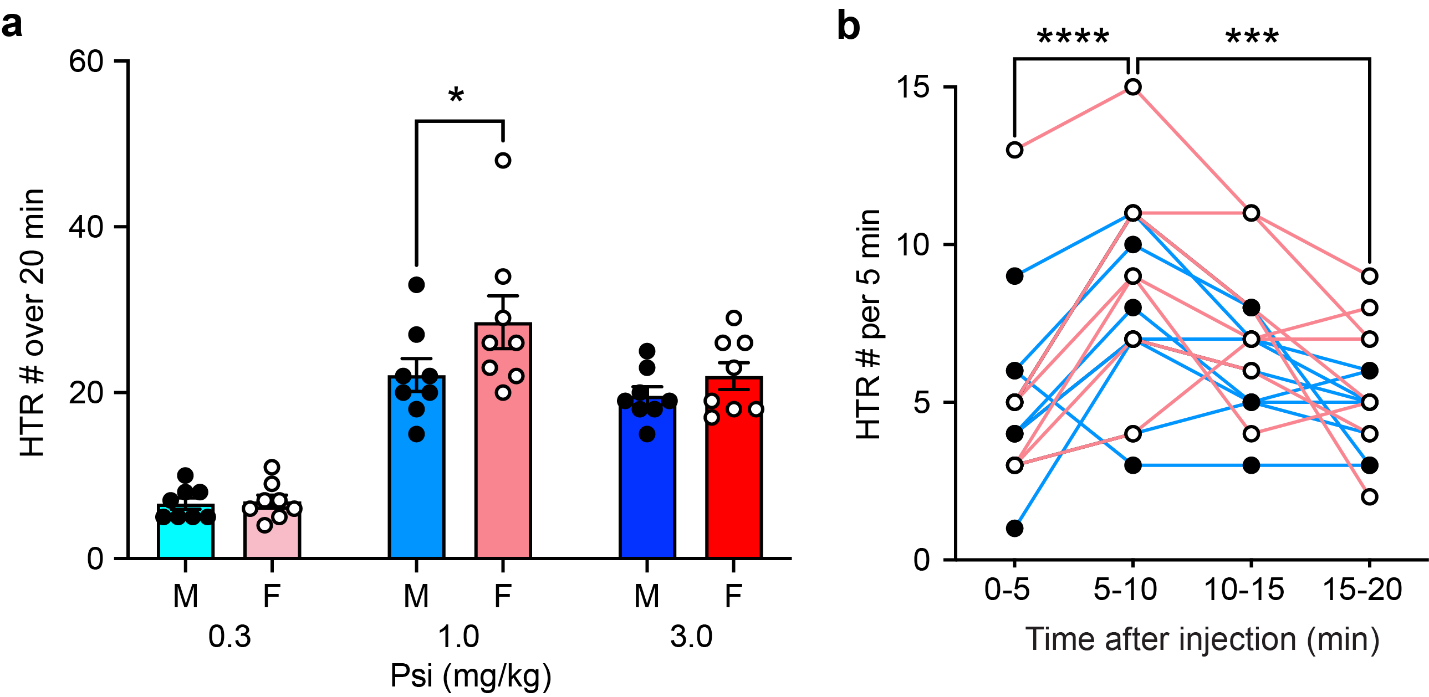

**Extended Data Figure 1.** Sex difference and time course of psilocybin-induced HTRs. **a**, HTR counts over 20 min after administration of different doses of psilocybin (Psi) in male and female mice. Two-way ANOVA followed by Šidák’s multiple comparisons test. Main effect of sex *F*(1,42) = 4.265, *p* = 0.0451; main effect of dosage *F*(2,42) = 59.25, *p* < 0.0001; interaction *F*(2,42) = 1.528, *p* = 0.2288. *n* = 8 males and 8 females per group. **b**, HTR counts after 1 mg/kg psilocybin administration counted in 5-min bins. Repeated measures one-way ANOVA *F*(3,45) = 11.53, *p* < 0.0001, followed by Tukey’s multiple comparisons test. *n* = 16 mice. Males are shown as filled circles and females as open circles. **p* < 0.05, ****p* < 0.001, *****p* < 0.0001.

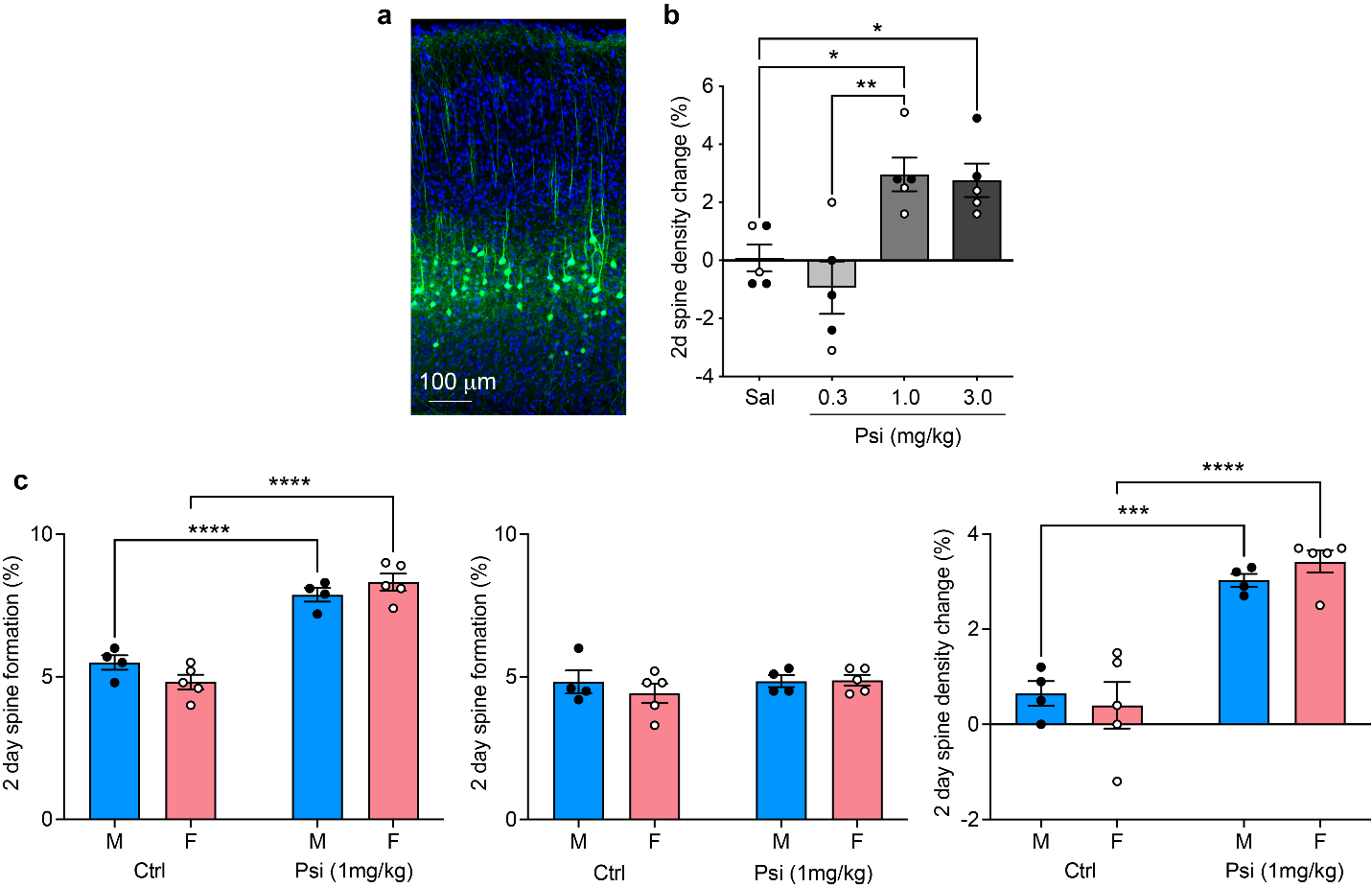

**Extended Data Figure 2.** Sex difference in spine dynamics in response to psilocybin treatment. **a**, Example of a coronal section of a *Thy1*-GFP-M mouse brain showing GFP+ neurons predominantly in cortical L5. **b**, 2d spine density changes in response to different doses of psilocybin. One-way ANOVA *F*(3,16) = 8.907, *p* = 0.0011, followed by Tukey’s multiple comparisons test. Sal: saline; Psi: psilocybin. *n* = 5 mice per dosage. **c**, 2d spine dynamics are comparable between male and female mice. Two-way ANOVA followed by uncorrected Fisher’s LSD test. Formation: main effect of sex *F*(1,14) = 0.1908, *p* = 0.6690, main effect of treatment *F*(1,14) = 119.2, *p* < 0.0001, interaction *F*(1,14) = 4.372, *p* = 0.0553. Elimination: main effect of sex *F*(1,14) = 0.3982, *p* = 0.5382, main effect of treatment *F*(1,14) = 0.6661, *p* = 0.4281, interaction *F*(1,14) = 0.5358, *p* = 0.4762. Spine density changes: main effect of sex *F*(1,14) = 0.04778, *p* = 0.8301, main effect of treatment *F*(1,14) = 66.14, *p* < 0.0001, interaction *F*(1,14) = 0.9454, *p* = 0.3474. *n* = 4 males and 5 females. Males are shown as filled circles and females as open circles. **p* < 0.05, ***p* < 0.01, ****p* < 0.001, *****p* < 0.0001.

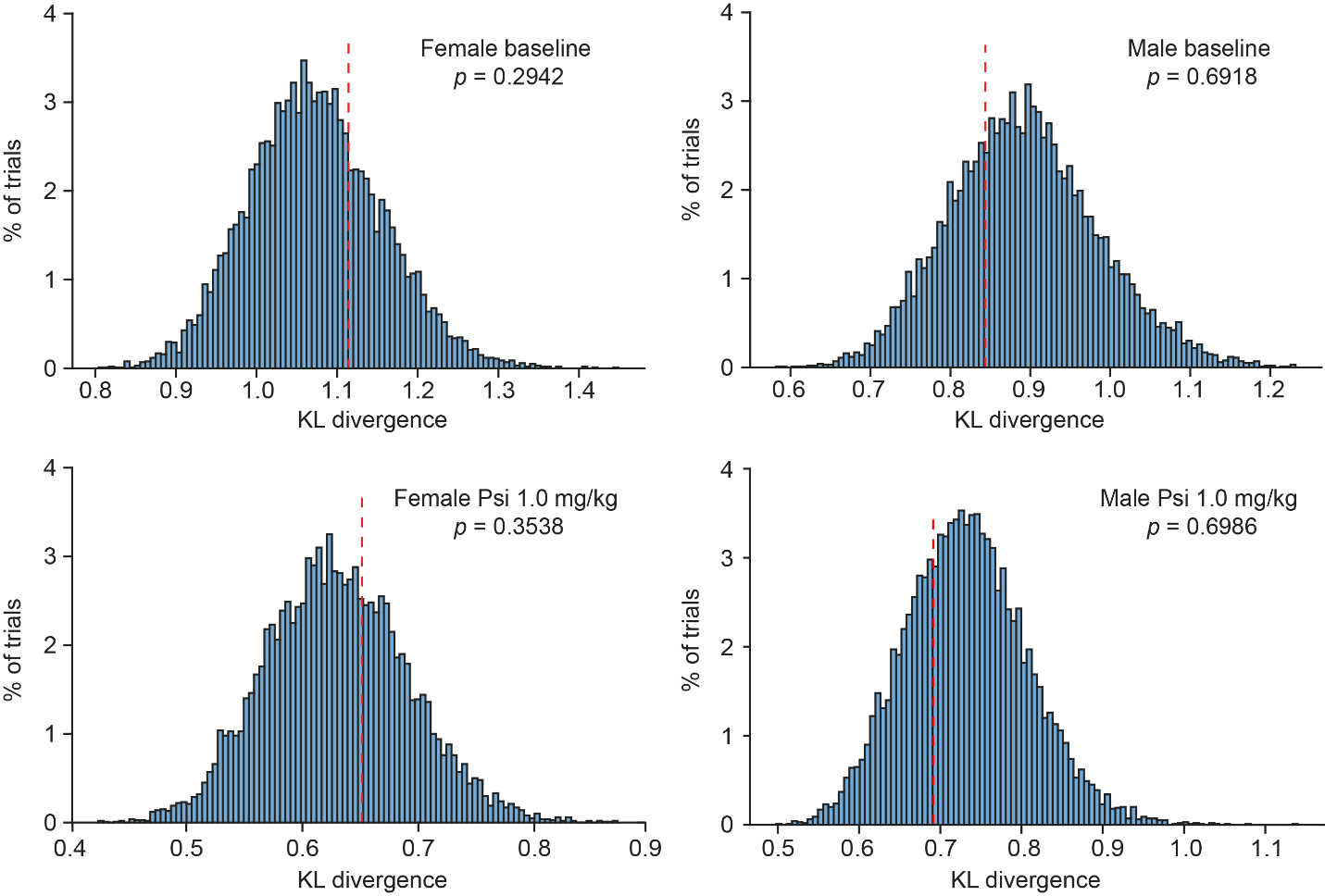

**Extended Data Figure 3.** Dendrite-level analysis of the uniformity of spine formation using a permutation test. The null model of uniform spine formation postulates that the probability of a new spine being formed on a dendrite is proportional to the initial spine count on that dendrite. In each trial of simulation, new spines are randomly and independently assigned to dendrites according to the null distribution. Because the number of spines assigned to each dendrite must be an integer, the simulated distribution fluctuates around the null distribution, and this deviation from uniformity is measured by the Kullback-Leibler (KL) divergence between the simulated and the null distribution. Each histogram shows the distribution of KL divergences of 10,000 trials of simulation. The red line marks the KL divergence between the empirical distribution (*i.e.*, experimentally observed dendrite-level spine formation) and the null distribution. The *p*-value indicates the percentage of simulated KL divergence values larger than the empirical value.

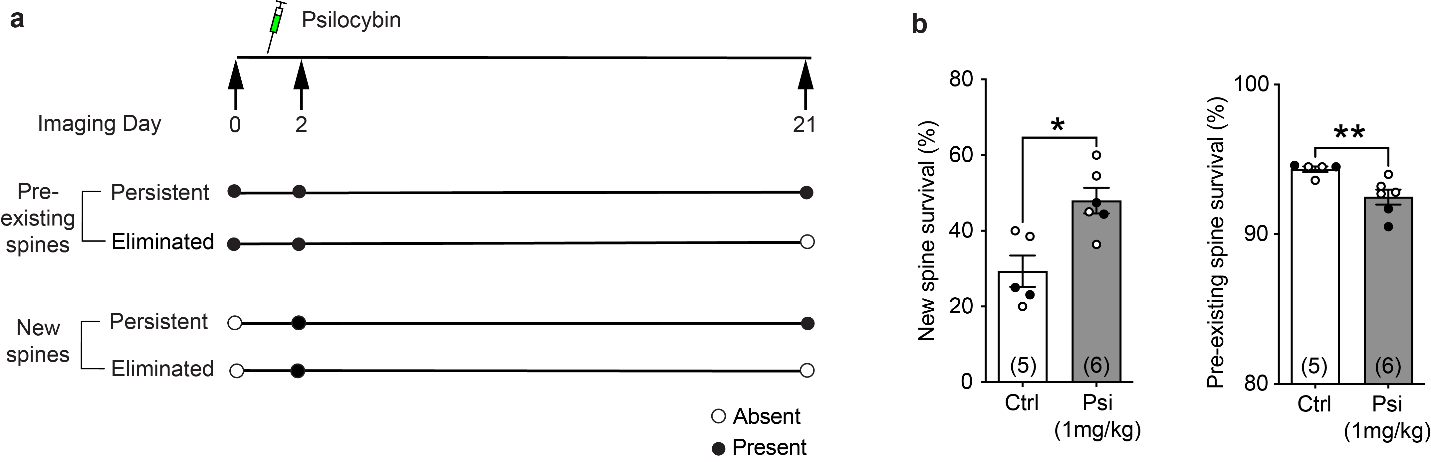

**Extended Data Figure 4.** Spine survival over 21d under baseline vs 1 mg/kg psilocybin treatment. **a**, Timeline of psilocybin administration and 2P imaging over 21d. **b**, 21d survival rate in control (Ctrl) vs psilocybin-treated (Psi) mice. Mann-Whitney test, new spines *U* = 2, *p* = 0.0173; pre-existing spines *U* = 1, *p* = 0.0087. *n* = # mice. Males are shown as filled circles and females as open circles. **p* < 0.05, ***p* < 0.01.

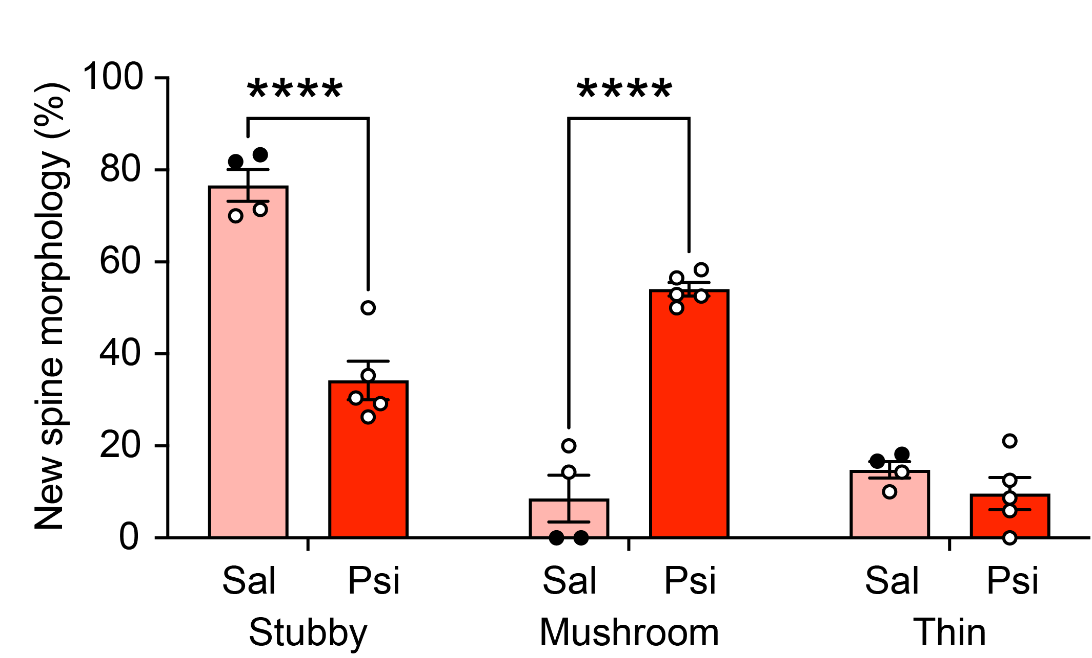

**Extended Data Figure 5**. Distribution of new spines across morphological categories in post-UMS mice. Two-way repeated measures ANOVA followed by Šidák’s multiple comparisons test. Main effect of morphology *F*(2,14) = 51.91, *p* < 0.0001; main effect of treatment *F*(1,7) = 2.003, *p* = 0.1999; interaction *F*(2,14) = 53.83, *p* < 0.0001. Sal: saline; Psi: 1 mg/kg psilocybin. *n* = 4 (Sal) and 5 (Psi) mice. Males are shown as filled circles and females as open circles. *****p* < 0.0001.

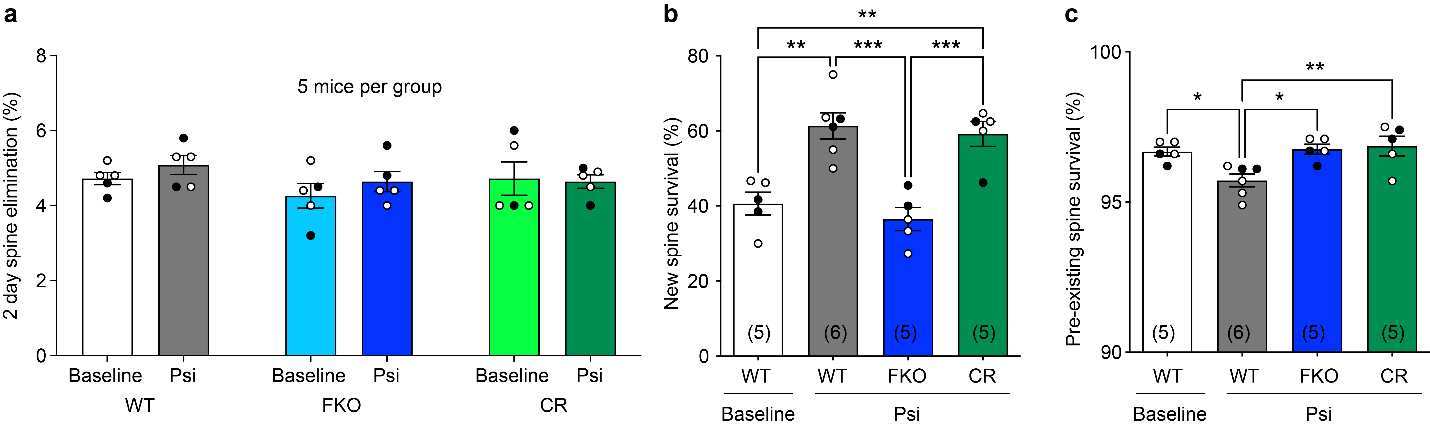

**Extended Data Figure 6.** Spine elimination and survival in WT, FKO, and CR mice with and without psilocybin treatment. **a**, 2d spine elimination at the baseline and after 1 mg/kg psilocybin treatment (Psi). Two-way ANOVA, main effect of treatment *F*(1,24) = 0.866, *p* = 0.3613; main effect of genotype *F*(2,24) = 1.208, *p* = 0.3163; interaction *F*(2,24) = 0.4032, *p* = 0.6726. **b**, 7d survival rate of new spines. One-way ANOVA *F*(3,17) = 14.99, *p* < 0.0001, followed by Tukey’s multiple comparisons test. **c**, 7d survival rate of pre-existing spines. One-way ANOVA *F*(3,17) = 5.999, *p* = 0.0056, followed by Tukey’s multiple comparisons test. *n* = # mice. Males are shown as filled circles and females as open circles. **p* < 0.05, ***p* < 0.01, ****p* < 0.001.

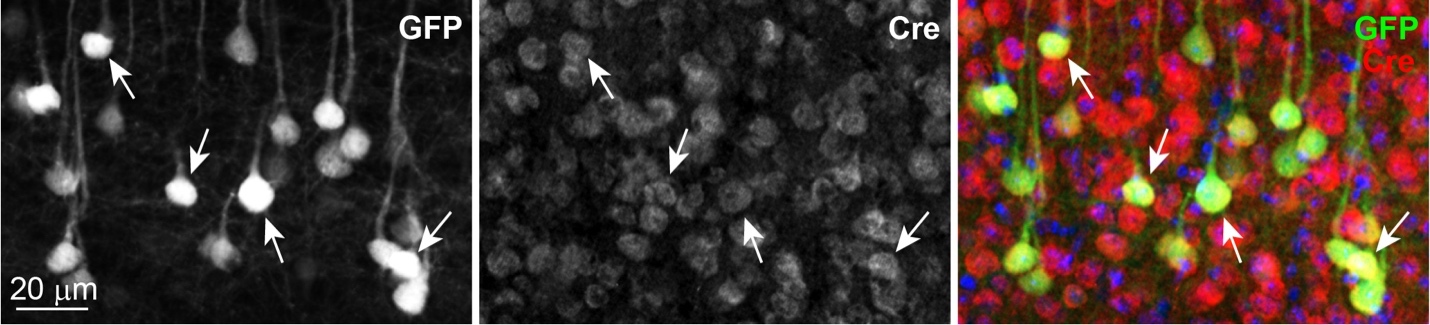

**Extended Data Figure 7.** Immunohistochemical labeling showing that the majority of GFP+ cells are also Cre+ in a CKO mouse.
